# Cognitive vulnerability to sleep deprivation is robustly associated with two dynamic connectivity states

**DOI:** 10.1101/509745

**Authors:** James Teng, Ju Lynn Ong, Amiya Patanaik, Jesisca Tandi, Juan Helen Zhou, Michael W.L. Chee, Julian Lim

**Affiliations:** Centre for Cognitive Neuroscience, Duke-NUS Medical School, Singapore, Singapore

**Keywords:** Sleep deprivation, individual differences, arousal, fMRI, dynamic functional connectivity, reproducibility

## Abstract

Robustly linking dynamic functional connectivity (DFC) states to behaviour is an important goal of the fledgling research using these methods. Previously, using a sliding window approach, we identified two dynamic connectivity states (DCS) linked to arousal. Here, in an independent dataset, 32 healthy participants underwent two sets of resting-state functional magnetic resonance imaging (fMRI) scans, once in a well-rested state and once after a single night of total sleep deprivation. Using a temporal differencing method, DFC and clustering analysis on the resting state fMRI data revealed five centroids that were highly correlated with those found in previous work, including the two states associated with high and low arousal. Individual differences in cognitive vulnerability to sleep deprivation were measured using changes in Psychomotor Vigilance Test (PVT) performance (lapses and median reaction speed), Changes in the percentage of time spent in the arousal states from the well-rested to the sleep-deprived condition specifically were correlated with declines in PVT performance. Our results provide good evidence of the validity and reproducibility of DFC measures, particularly with regard to measuring arousal and attention, and are an encouraging base from which to build a chronnectome mapping DCS to cognition.

---

Dynamic functional connectivity (DFC) analysis of neuroimaging data is increasingly being used to study how inter-region connectivity strength and network configurations evolve over time (Hutchison et al. 2013). One common approach in DFC is to employ clustering analysis on all windows within a connectivity time series (Allen et al. 2014) to obtain recurring patterns known as dynamic connectivity states (DCSs). There is now some evidence that DCSs contain information that is of behavioural significance. For instance, on a coarse level, certain DCS have been associated with wakefulness and sleep (Haimovici et al. 2017, Damaraju et al. 2018), motivating a more detailed search for states related to other cognitive domains.

In the short time since the first reports of time-varying connectivity, a plethora of different approaches have been proposed to derive DFC estimates, each with their own theoretical underpinnings. These approaches vary along three major dimensions: 1) transformations (if any) applied to the data, 2) the nature of the function that quantifies the relationships between time series, and 3) the weighting vectors applied to the relational computation (Thompson and Fransson 2018).

Moreover, within each analysis strategy, there are a considerable number of parameters that can be tuned (e.g. window size for analyses involving moving averages, component selection for ICA approaches). This heterogeneity in existing methods can lead to challenges in the interpretation and comparison of findings, particularly when researchers make claims that particular DCS are linked to specific cognitive or behavioural states.

Several reports have been published on the reliability of DFC estimates, reaching the general conclusion that reliability is good for summary statistics (e.g. average connectivity, percentage of occurrence) and connectivity features (Abrol et al. 2017, Choe et al. 2017, Smith et al. 2018), but relatively lower for derived measures such as dwell time and transitions. However, it is typically these latter measures that are used in a search for DFC-behaviour relationships. It thus follows that a system employed to test the robustness of DFC-behaviour associations must be anchored by a highly reliable behavioural phenomenon.

Accordingly, the properties of sustained attention and arousal as they are affected by sleep deprivation (SD) make this paradigm an attractive test bed to harmonize results across connectivity experiments. Acute sleep deprivation (SD) causes reversible but serious decrements in sustained attention (Lim and Dinges 2010). Individual differences in these impairments are stable over time (Leproult et al. 2003, Van Dongen et al. 2004), and this trait-like nature makes them especially amendable to reproducibility studies. Indeed, changes in BOLD activation in frontoparietal regions are also reproducible across two nights of total SD (Lim et al. 2007), paralleling the findings from behavioural work.

Using static, or time-averaged connectivity analysis, it has been shown that the brain connectome is less integrated and segregated in SD (Yeo et al. 2015), and that anti-correlations stemming from the default-mode network (DMN) are particularly implicated in this global change (Samann et al. 2010, De Havas et al. 2012). Using DFC analysis, Xu et al. (2018) showed that large shifts in the proportions of dwell times and the transition probability matrix occur in resting-state data after 36 h of total SD. Separately, two prior DFC studies have reported DCSs associated with sustained attention (the high arousal state (HAS) and low arousal state (LAS)), showing that these DCSs in the sleep-deprived state are associated with temporal fluctuations in vigilance at rest and in an auditory vigilance task condition (Wang et al. 2016), and can be used to expose vulnerability to SD while individuals are still in a well-rested state (Patanaik et al. 2018).

In the current study, we exploited SD as a tool to test the robustness of the HAS/LAS behavioral associations across datasets analysis methods (i.e. in comparison with our previous published findings).Specifically, we sought to elaborate on our previous findings by directly investigating SD-related individual differences in DCSs, and vigilant attention. To achieve this, we collected resting-state fMRI data at baseline and after 24 hours of total sleep deprivation in a group of 32 healthy young adults. Our primary hypothesis was that following SD, we would observe reductions in a composite measure calculated from proportions of two DCSs previously shown to index high and low arousal states (Patanaik et al. 2018), and that this decrement would correlate with changes in sustained attention. A third state previously associated with trait mindfulness (Lim et al. 2018), and two other unnamed (but reproducible) states were also tested to demonstrate the specificity of this effect. Critically, we used a different method of DFC computation (multiplication of temporal differences) than in our original reports on DCS relationship with arousal for all these analyses. Finally, we tested two other variables known to relate to arousal – global signal variability and head motion – to assess their independent contribution beyond that of the DCSs.

## Materials and Methods

### Participants

32 participants were recruited from the National University of Singapore through online advertising and word-of-mouth as part of a larger study to investigate the effects of sleep deprivation. Data from two of these participants was discarded after the first-pass connectivity analysis (see below), resulting in a total of 30 participants (15 males; mean age (sd) = 23 (3.59)). All participants were screened for right-handedness (Oldfield 1971) and normal or corrected-to-normal vision, and to ensure they had no history of long-term physical or psychological disorders. This study was approved by the National University of Singapore Institutional Review Board, and all participants provided written informed consent.

### Psychomotor vigilance test

The Psychomotor Vigilance Test (PVT, Dinges 1995) is a sensitive assay of sustained attention under conditions of fatigue and sleep loss (Lim and Dinges 2008). In the task, participants monitor a rectangular box in the center of a screen and respond as quickly as possible to the appearance of a millisecond counter. The number of lapses (reaction time > 500 ms), and response speed (RSp; reciprocal reaction time) on this test are robust markers of vigilance (Basner and Dinges 2011). PVT stimuli were presented using Psychtoolbox (Brainard, 1997) in MATLAB 2012a.

### Study protocol

Participants were invited to the lab for two counterbalanced sessions held approximately one week apart: a night of rested wakefulness (RW), where they were required to adhere to a strict 9 h sleep opportunity, and a night of total sleep deprivation (SD). They were required to maintain a consistent sleep-wake schedule (to sleep before 0030 and wake up before 0900, with approximately 6.5 to 9 hr sleep opportunity a night) for approximately one week before each of these testing nights; this was verified by wrist actigraphy (Actiwatch 2, Philips Respironics Inc., Pittsburgh, PA) worn on the non-dominant hand for the duration of the study.

Participants arrived at the lab at approximately 1900 on both experimental nights, and performed a battery of cognitive tasks (results not reported). Participants in the SD session were then given computerised tasks every hour to help them stay awake with the PVT given every other hour. Participants in the RW session were given a 9 hr sleep opportunity. A final PVT was performed in the morning before fMRI scans at 0730 for RW sessions after participants were awakened and given 30 minutes to recover from sleep inertia, and at 0600 for SD sessions. This final PVT, compared with baseline performance on the RW night, was used to measure the effect of SD on vigilance.

### fMRI acquisition

Functional MRI scans were collected on a 3-Tesla Siemens PrismaFit system (Siemens, Erlangen, Germany) using an interleaved gradient echo-planar imaging sequence (TR = 2000 ms, TE = 30 ms, FA = 90°, FoV = 192 × 192 mm, voxel size = 3 × 3 × 3 mm). Thirty-six oblique axial slices were obtained, and 180 volumes were collected for each. Concurrent eye videos were acquired using an MR compatible camera (NNL EyeTracking Camera, NordicNeuroLab) placed over the right eye. Two runs of 6 min (180 TR) eyes-open resting state (RS) scan were collected at the beginning of an approximately hour-long fMRI scanning session. During the RS scan, participants were instructed to remain still and keep their eyes open. Pre-recorded wake-up calls (e.g., “Open your eyes.”) were delivered whenever participants closed their eyes for more than 10 s. We have previously used this procedure to ensure that participants remain awake during RS scans under conditions of high sleep pressure (Yeo et al. 2015), as this can confound the results of connectivity analysis (Tagliazucchi and Laufs 2014). High-resolution structural images were collected using an MPRAGE sequence (TR = 2300 ms, TI = 900 ms, FA = 8°, voxel size = 1 × 1 × 1 mm, FOV = 256 × 240 mm, 192 slices).

### RS-fMRI analysis

RS scans were preprocessed in accordance to the previously described procedure in Yeo et al. (2015). Briefly, preprocessing steps involved discarding the first four frames of each run, slice time correction, head-motion correction, linear trend removal, and low-pass temporal filtering. White matter, ventricular signals, and motion parameters and their associated derivatives were regressed out, and functional data of individual subjects were then projected onto MNI-152 space, downsampled to 2 mm voxels and then smoothed with a 6-mm full width half maximum kernel. Global signal regression (GSR) was carried out as a part of the preprocessing pipeline, and global signal power, or the standard deviation of the average percentage change in the signal time course of the whole brain (Wong et al. 2013), was calculated. Volumes having framewise displacement (FD) > 0.2 mm or DVARS (Power et al. 2012) >5% were marked as high motion. As dynamic functional connectivity analysis was the intended analysis, motion scrubbing was not conducted. Instead, one volume before and two volumes after each high motion volume were also marked, and these frames were interpolated from surrounding data. No subject was excluded from the analysis for having more than 50% of total volumes marked as high motion.

### Dynamic functional connectivity analysis

DFC analysis was performed using the multiplication of temporal derivatives (MTD) method described by Shine et al. (2015), as this method has been shown to be more sensitive to dynamic alterations in connectivity structure compared to traditional sliding-window methods (Allen et al., 2014). The coupling between each pairwise set of 114 ROIs (Yeo et al. 2011) was estimated by multiplying the first-derivatives of the averaged BOLD time series. Connectivity at each time point was then estimated by computing a simple moving average of the MTD time course using the recommended window size of 7 TRs, for a total of 168 coupling matrices per participant, each containing 6441 (114 × 113/2) unique coupling values.

Coupling matrices were than concatenated across the 30 participants and k-means clustering was performed to classify each matrix using squared Euclidean distance as the cost function. We elected to use a k = 5 solution, as recent work using a large (N = 7,500) resting-state dataset suggests that this is an optimal number of clusters (Abrol et al. 2017). Our initial analysis using this approach revealed two artefactual states consisting of only positive values that were unique to two individuals. Data from these participants was removed, and matrices from the remaining 30 participants re-clustered. To confirm that our centroids were consistent with those obtained from a previous analysis reported by our group (Lim et al. 2018), we performed Spearman’s correlations between the two sets of centroids. We also calculated the proportion of the run spent in each DCS, as well as the number of percentage transitions, defined as the proportion of frames in the time course that differ in state classification from one time point to the next.

### Arousal Index

Previous studies have suggested a moderate relationship between arousal and two particular DCSs, the high and low arousal state (HAS and LAS respectively; Patanaik et al. 2018). Following the methods reported here, an arousal index (AI) was calculated as a summary measure of the proportions of time spent in the HAS (T_HAS_) and LAS (T_LAS_), using the formula AI = 1 + T_HAS_ – T_LAS_.

### Statistical analysis

Statistical analysis was performed using SPSS (version 25, Armonk, NY: IBM Corp), and statistical significance for all analysis was set at α = 0.05. Dependent variables of interest from the RW and SD nights were compared using independent-samples t*-*tests, specifically: PVT results (lapses and RSp), proportion of time spent in DCSs, percentage state transitions, arousal index, global signal, and head motion.

A change score was then calculated for each of these variables (e.g. Δlapse = lapse on SD nights – lapse on RW nights) and Pearson’s correlation was used to assess the linear relationship between the change scores of PVT performance with proportion of DCSs, percentage state transitions, AI, and head motion. To control for the effects of global signal, we entered this variable into a multiple linear regression together with PVT performance and AI.

## Results

### Behavioural measures

To establish that the night of total sleep deprivation negatively affected vigilance, we conducted paired-samples t-tests on lapses (reaction time > 500 ms) and reaction speed (RSp) on the PVT. As expected, participants responded faster in RW compared with SD (Fig 1A. RSp in RW: mean(sd) = 3.12 (0.298); RSp in SD: mean(sd) = 2.79 (0.287); *t_29_* = 4.75, *p* < 10^−5^). Similarly, fewer lapses occurred in RW compared to SD (Fig 1B. RW: mean(sd) = 3.57 (3.94); SD: mean(sd) = 13.6 (10.28); *t_29_* = 5.53, *p* < 10^−6^). Due to the non-normal distribution of lapses, lapses were normalised (Dinges et al. 1997; √n + √(n+1)) prior to subsequent analyses.

**Figure 1.**
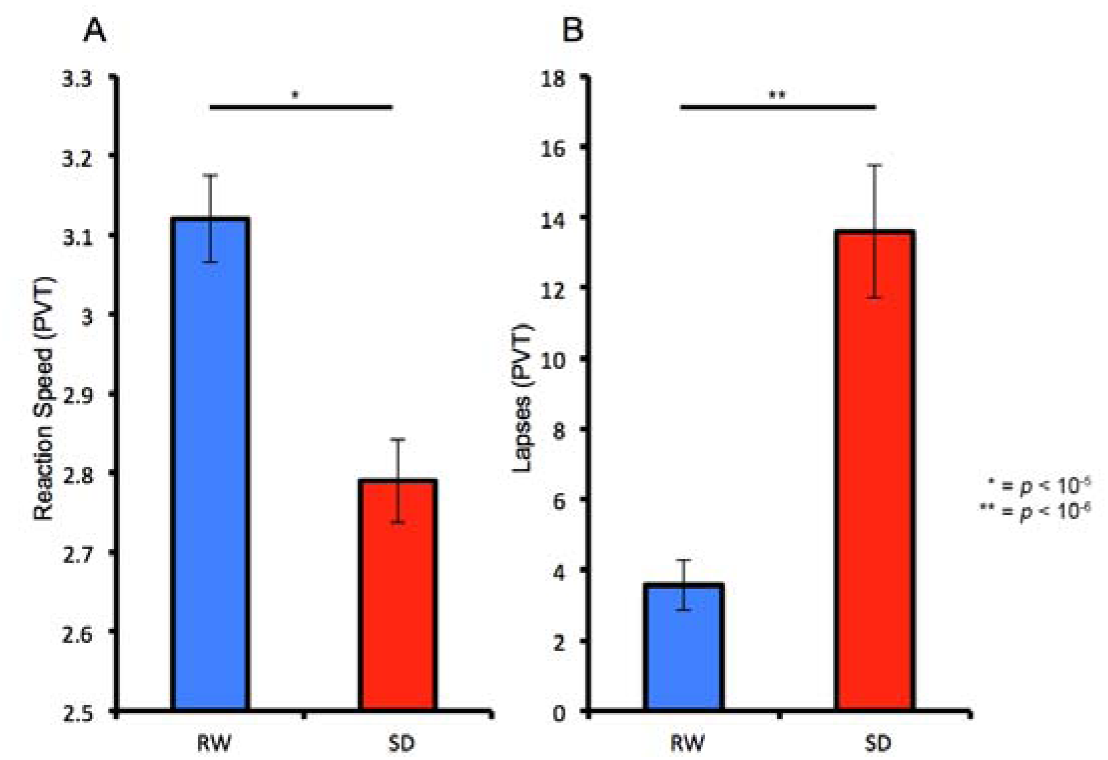
One night of total sleep deprivation (SD) resulted in A) slower responding, and B) more lapses (reaction times > 500 ms) on a 10-minute Psychomotor Vigilance Test (PVT) compared with rested wakefulness (RW)

### Reproducibility of dynamic connectivity states

Due to the different conditions and methodologies under which the DCSs were obtained, a comparison between the current DCSs (Fig 2A) and those found in prior work was performed using Spearman’s rank-order correlation to ensure congruence. Consistent with prior findings, three dynamic connectivity states (DCSs) that resembled the, LAS, HAS, and the task-ready state (TRS) were reproduced (Wang et al. 2016, Lim et al. 2018); these states were highly correlated with centroids obtained using the MTD method in Lim et al. (2018); r_s-TRS_ = 0.89, r_s-LAS_ = 0.90, r_s-HAS_ = 0.91) and moderately correlated with centroids obtained using the sliding-window approach (Patanaik et al. 2018): r_s-TRS_ = 0.77, r_s-LAS_ = 0.60, r_s-HAS_ = 0.74). The characteristics of the named states are as follows:

A. The LAS features lower within-network correlations and relatively small anti-correlations between task-positive networks (dorsal attention network (DAN), ventral attention/salience network (VAN), executive control network (ECN)) and the default-mode network (DMN).
B. The HAS features higher within-network connectivity in the DMN, ECN, VAN, and DAN, as well as higher between-network connectivity between DMN and ECN, and VAN and DAN. Greater anti-correlations were also found between the DMN and DAN/VAN.
C. The TRS features stronger within-network correlations in the DMN and the VAN, and larger anti-correlations between task-positive networks.

**Figure 2.**
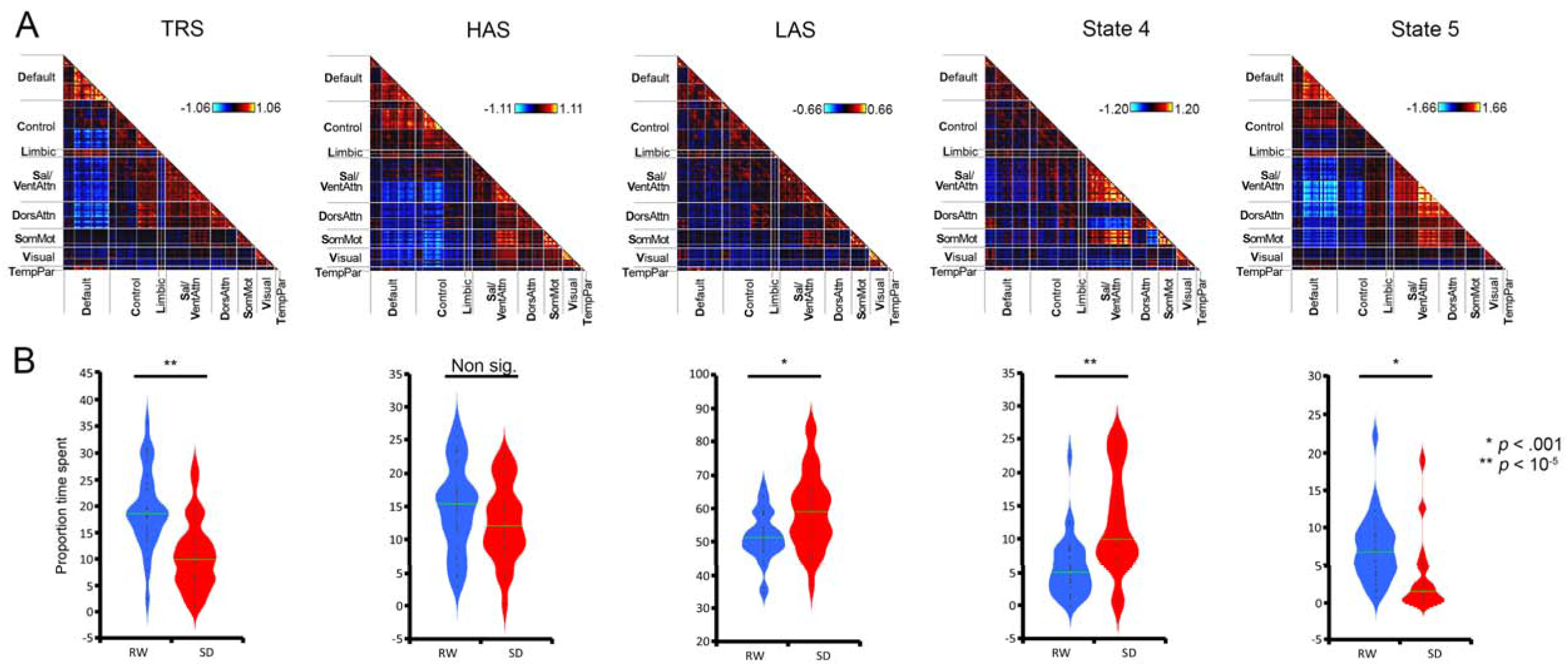
A) Connectivity centroids derived from k-means clustering across all connectivity matrices across rested wakefulness (RW) and sleep deprivation (SD). Previously described states are the task-ready state (TRS), the high arousal state (HAS), and the low arousal state (LAS). B) Violin plots showing distributions of proportions of time spent in each state in RW and SD. Significant increases were observed in the time spent in LAS and State 4, and significant decreases seen in time spent in TRS and State 5.

Of note, while the HAS and the TRS have similar features, the TRS has previously been associated with trait mindfulness (Lim et al. 2018), while the HAS and LAS have been associated with fluctuations in vigilance (Wang et al. 2016), and are predictive of vulnerability to sleep restriction (Patanaik et al. 2018). The remaining 2 states in the k = 5 solution are also reproducible across datasets (Lim et al. (2018): r_s-state4_ = 0.74, r_s-state5_ = 0.88; Patanaik et al. (2018): r_s-state4_ = 0.66, r_s-state5_ = 0.80), but have not yet been ascribed any functional significance.

### Change in dynamic functional connectivity following sleep deprivation

For comparability with previous reports, AI in RW and SD nights was calculated. Paired-samples t-tests showed that AI was higher in RW scans compared to SD (t_29_ = 2.74, *p* = .010). We computed the total time spent in each of the five states across RW and SD (Fig 2B). Of the previously named DCSs, participants spent significantly less time in LAS on RW than SD nights (t_29_ = 3.16, *p* = .0039) and more time in TRS on RW than SD nights (t_29_ = 5.32, *p* < 10^−5^). Counter to our expectations, no significant decrease was found in time spent in HAS on the SD night (t_29_ = 1.43, *p* = .16). Of the unnamed DCSs, participants showed an increase in State 4 after RW (t_29_ = 4.53, *p* < 10^−5^), while less time was spent in State 5 following SD (t_29_ = 2.92, *p* = .0067).

The percentage of overall transitions also differed between the nights, with participants transitioning between states more often in the RW nights than SD nights (Fig 3A; t_29_ = 3.46, *p* = .002).

**Figure 3:**
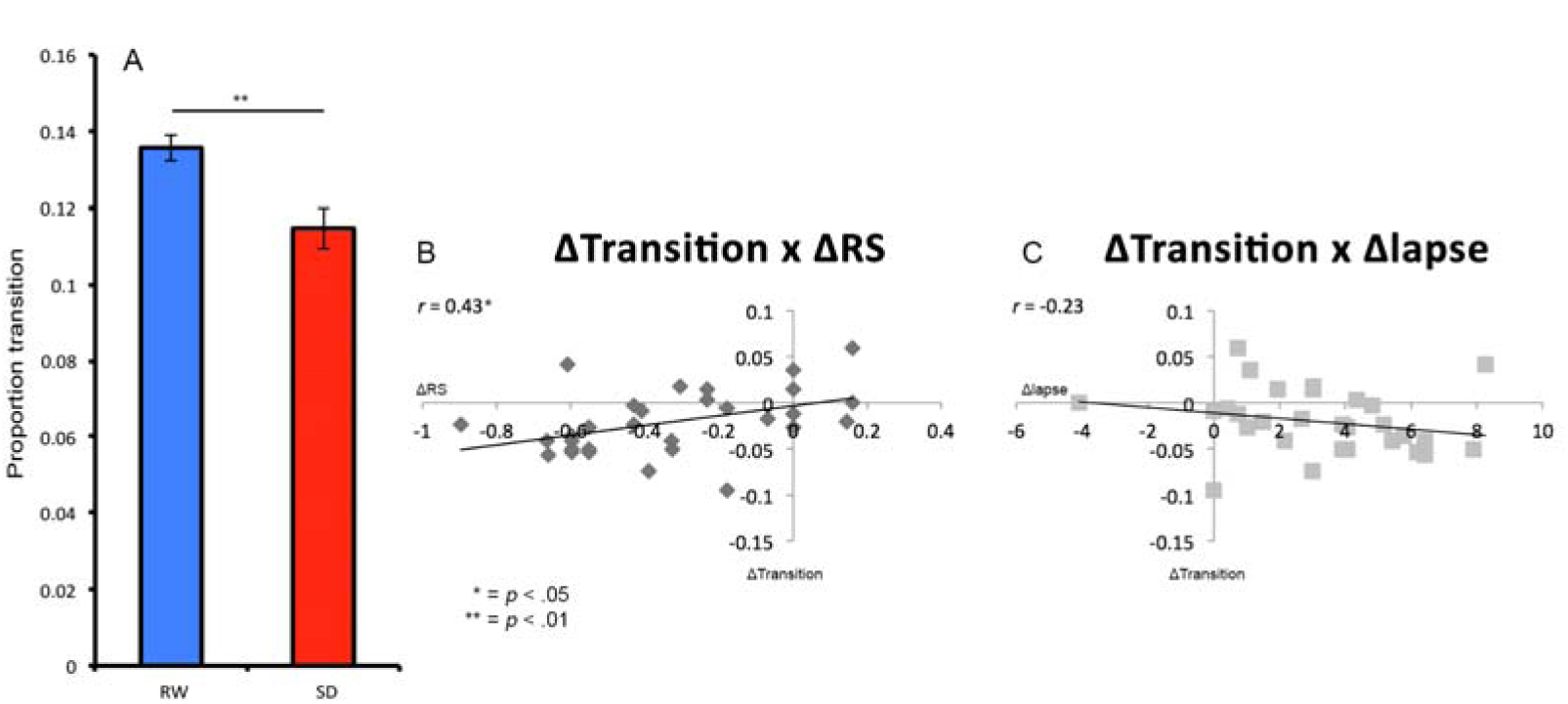
A) Transitions between dynamic connectivity states (DFC) decrease from rested wakefulness (RW) to sleep deprivation (SD). Values reflect the proportion of volumes when a transition occurred. The change in transition proportion across state (RW –SD) correlates with B) change in reaction speed (ΔRS), and C) change in normalised lapses (Δlapse).

### Connectivity-behaviour relationships

Central to the current investigation is the relationship between the effect of sleep deprivation on DFC and on vigilance. To examine this, we correlated the change scores for PVT performance (ΔRSp and Δlapse) with those for the proportion of time spent in each DCS. Correlations with the difference in the percentage transitions and AI were computed as well to interrogate the effects of more global DFC variables.

Change in percentage LAS (ΔLAS) across sleep conditions correlated significantly with both ΔRSp (Fig 4A; *r* = −0.64, *p* = < .0001), and Δlapse (Fig 4B; *r* = 0.43, *p* = .018). Change in percentage HAS (ΔHAS) were also significantly correlated with both ΔRSp (Fig 4E; *r* = 0.43, *p* = .019), and Δlapse (Fig 4F; *r* = −0.39, *p* = .033), even though the proportion of time spent in HAS across RW and SD were not significantly different. Changes in AI (ΔAI) were also significantly correlated to both ΔRSp (Fig 5A; *r* = 0.6090, *p* < .0004) and Δlapse (Fig 5B; *r* = −0.4550, *p* = .012). In contrast, change in percentage TRS (ΔTRS) did not correlate with either ΔRSp (Fig 4C; *r* = 0.35, *p* = .056), or Δlapse (Fig 4D; *r* = −0.22, *p* = .25). Similarly, we did not find correlations between changes in percentage State4 (ΔState4) and percentage State5 (ΔState5) and either PVT measure (all *p* > .05).

**Figure 4.**
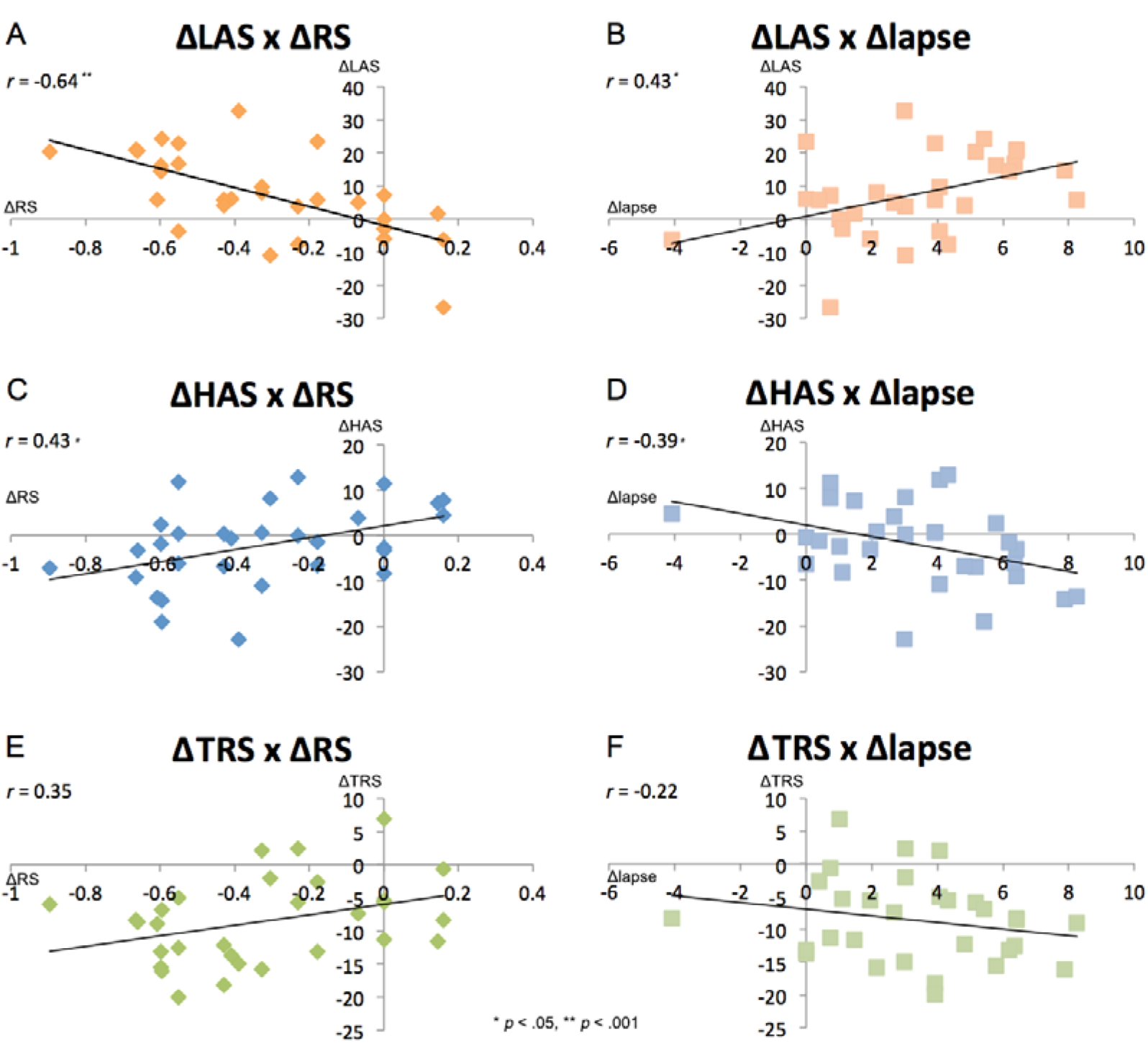
Correlations between dynamic connectivity states (low arousal state (LAS), high arousal state (HAS), task-ready state (TRS)) and behaviour (response speed (RS) and normalised lapses). A-D) ΔLAS and ΔHAS are significantly correlated with both the change in both behavioural metrics across state, while E-F) ΔTRS was not correlated with behavioural change.

**Figure 5.**
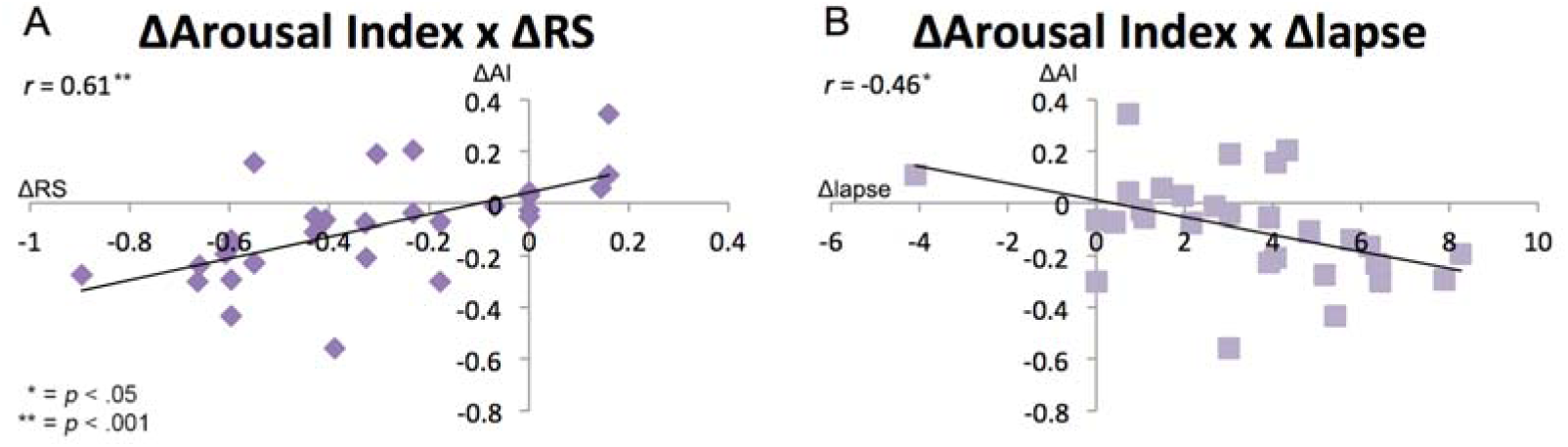
Change in an arousal index (AI = 1 + T_HAS_ – T_LAS_) across state is significantly correlated with A) changes in response speed and B) normalised lapses.

In addition, change in percentage transitions (ΔTransitions) correlated with ΔRSp (Fig 3B; *r* = 0.43, *p* = .017), but not Δlapse (Fig 3C; *r* = −0.23, *p* = .22).

### Changes in global signal

As GS has previously been related to vigilance (Wong et al. 2013), we analysed this variable to assess its independent contribution to this outcome variable. As expected, GS was significantly lower in RW than SD (t_29_ = 5.86, *p* < 10^−6^) (Yeo et al. 2015, Nilsonne et al. 2017). To further interrogate the relationship between GS and vigilance directly, we performed correlation analysis between changes in GS (ΔGS) and the two PVT measures. We found that while the ΔGS correlation with ΔRSp was below the threshold of statistical significance (Fig 6A; *r* = - 0.32, *p* = .082), there was a significant correlation between ΔGS and Δlapse (Fig 6B; *r* = 0.42, *p* = .020).

**Figure 6.**
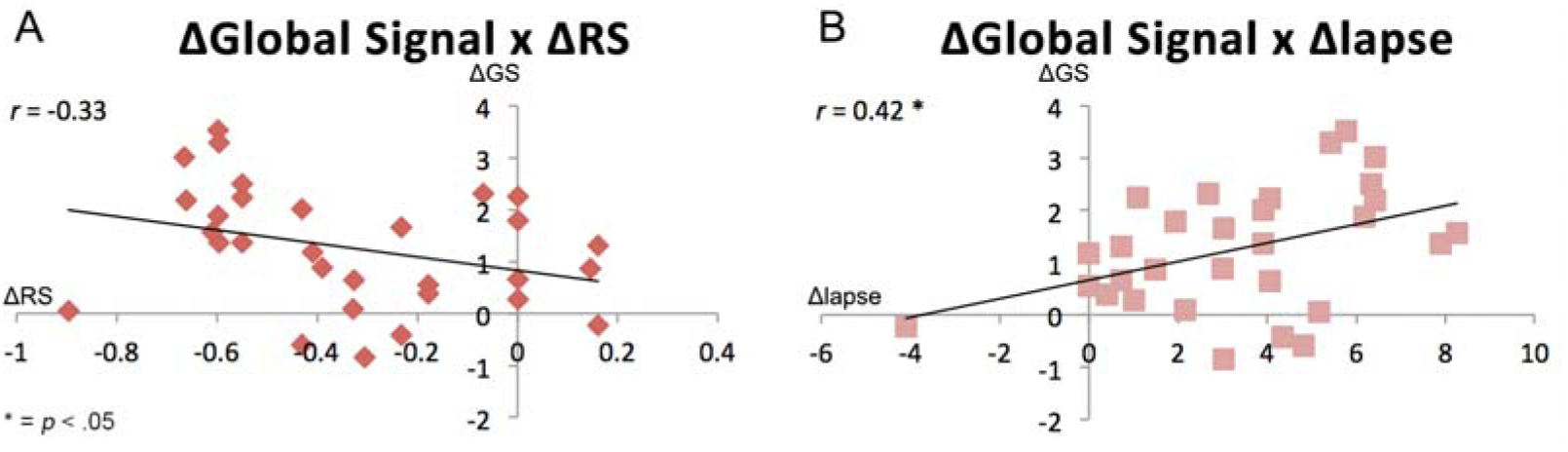
Changes in global signal across state are A) not correlated with response speed, but B) significantly correlated with normalised lapses.

Since ΔGS was an independent predictor of Δlapse, we entered both ΔAI and ΔGS into a multiple linear regression to assess their independent contributions to this variable. We found that while with both ΔGS and ΔAI predicted Δlapse (R^2^ = .29, F(2, 29) = 5.63, *p* = .009), ΔAI significantly contributed to the model (β = −4.98, *p* = .046) but ΔGS did not (β = .741, *p* = .079).

### Changes in head motion

Head motion is a significant source of noise in connectivity analysis (Power et al. 2012), and also typically increases following sleep deprivation. Accordingly, we compared head motion parameters between RW and SD, and found a trend to more movement in SD (DVARS_RW_: mean(sd) = 23.29 (2.90); DVARS_SD_: mean(sd) = 24.71 (2.89); *t_29_* = 1.94, *p* = .062). We then examined the possible influence of head motion on the connectivity-behaviour correlations. Change scores of DVARS (ΔDVARS) across sleep conditions did not significantly correspond to both ΔRSp (*r* = .02, *p* =.93) and Δlapse (*r* = .008, *p* = .97).

When entered into a partial correlation, ΔDVARS did not significantly alter the relationship between ΔAI and ΔGS on PVT measures (ΔAI × ΔRSp: *p* = .93; ΔAI × Δlapse: *p* = .97; ΔGS × ΔRSp: *p* = .93; ΔGS × Δlapse: *p* = .97). This finding held when substituting framewise displacement for DVARS (all *p* > .05).

## Discussion

Finding robust links between dynamic functional connectivity and behaviour is an important on-going endeavour. Here we show that DCS centroids are reproducible across datasets and analysis methods, and further demonstrate that total sleep deprivation substantially alters the profile of these dynamic connectivity states (DCSs) in individuals. Importantly, these DCS changes are closely tied to behavioural performance, as measured by declines in sustained attention, the cognitive module that is most significantly affected by sleep loss (Lim et al. 2010), even after accounting separately for the contributions of global signal and head motion, both of which has been known to increase following SD. While decrements in vigilance were accompanied by decreases in the occurrence of highly integrated DCSs (e.g. HAS, TRS), and increased proportions of DCSs with low integration (e.g. LAS), only the two specific DCSs previously identified as “arousal” states (HAS/LAS) were correlated with behavioural change.

### DCSs are comparable across sliding window and temporal differencing methods

Since the first reports of DFC analyses using the sliding window Pearson’s correlation (SWPC; Handwerker et al. 2012, Allen et al. 2014), methodological advancements have created a range of options for investigators aiming to study connectivity fluctuations (Thompson and Fransson 2018). However, few studies to date have performed head-to-head comparisons of these methods, or assessed the robustness of findings across approaches. As such, methodological differences pose a challenge to comparisons between reports across the field.

In the current analysis, we used the multiplication of temporal differences method and compared the findings against our previous reports using SWPC. Encouragingly, we found a similar pattern of information with high spatial correlations between the two methods across multiple studies (Lim et al. 2016, Wang et al. 2016, Patanaik et al., 2018). MTD has been found to be less sensitive to low frequency drifts, due to the inherent nature of differencing acting as a high-pass filter, but also to have less signal-to-noise ratio (Ochab et al. 2019). As a result, the susceptibility of MTD to higher frequency signal (Shine et al. 2015) as compared to SWPC necessarily means that the properties of the connectivity information it contains are different. Nevertheless, the similarity of the spatial patters in the resultant centroids lead us to speculate that these specific state patterns might represent stable “attractor” states (van den Heuvel and Sporns 2011) that are indifferent to the frequency of brain information that is being sampled. This is a particularly important finding for those seeking to create a cohort-common, canonical chronnectome (Calhoun et al. 2014) to function as an atlas for the neuroimaging community.

### Sleep deprivation affects the profile of DCS occurrence

Having established the comparability of our named DCSs, we next examined the effect of total sleep deprivation (TSD) on the occurrence of those states. Following TSD, we found a change in four of the five chosen states: decreases in the TRS and state 5, and increases in the LAS and state 4. Prior work has found distinct patterns of time-averaged functional connectivity between sleep-deprived participants and those who received a full night of sleep (Yeo et al. 2015, Kaufmann et al. 2016), with greater connectivity magnitude apparent during RW relative to SD (Samann et al. 2010, De Havas et al. 2012, Yeo et al. 2015). Our findings are largely in line with these reports, as evidenced by the significant declines in the occurrence of states with strong DMN anti-correlations, while states with less DMN anti-correlations increased following TSD.

A recent study (Xu et al. 2018) using DFC analysis on a group of subjects sleep deprived for 36 hours found that DCSs linked specifically to RW and SD conditions occurred in different proportions across sleep conditions. In agreement with our findings, RW-associated states generally featured greater DMN anti-correlations. This correspondence of strong DMN anti-correlated states being more predominant during rested wakefulness was found despite a difference in methodology and the inclusion of global signal in the Xu et al. report.

A closer examination of state 4 in our analysis also revealed lower DMN anti-correlations similar to the LAS, but also high integration of the salience network – a state that also showed an increase in proportion following TSD. Due to the role of the salience network in stimulus detection and subsequent redirection of attention (Seeley et al. 2007), we propose that the increase in proportion of this DCS might represent overcompensation by the brain in an attempt to remain awake following TSD (Doran et al. 2001, Ong et al. 2013)

Of our named DCSs, the HAS was not significantly altered following TSD, even though it too displayed a characteristically strong DMN anti-correlations with task-positive areas. Prior work has suggested that these highly integrated organizations might represent a costly, but highly efficient network to which the brain may enter, either spontaneously or in response to task demands (Bullmore and Sporns 2012). The high spatial similarity between task-based and resting state DCSs (Wang et al. 2016) supports this. Indeed, across both sleep conditions, we observe that LAS occurs at more than twice the rate of HAS. We speculate from this null finding that a certain proportion of time must be spent in the metabolically costly HAS in order to sustain some level of wakefulness, even under conditions of high homeostatic sleep pressure. This is in line with our theory the HAS is a fundamental state that is essential for timely responding to exogenous stimuli.

### DFC states are an index of individual differences in SD vulnerability

The key finding of this study was that previously defined arousal-related DCSs were associated with individual differences in vigilance declines following SD. Specifically, we used an arousal index (AI; Patanaik et al. 2018) comprising the proportion of time spent in HAS and LAS (Wang et al. 2016) and found that decreases in AI were correlated with decrements in PVT performance. Correlations were also observed between behavioural changes and the individual constituents of AI.

To date, the strongest links between physiology/behaviour and DFC metrics have been made in the domains of arousal and vigilance. Connectivity fluctuations can serve as an index of wakefulness and sleep (Tagliazucchi and Laufs 2014, Haimovici et al. 2017), and can also track online arousal levels (Chang et al. 2016). Two particular studies directly motivated the investigation reported in this experiment. First, Wang et al. (2016) reported that spontaneous eye closures can be used as a proxy for arousal due to its long association with vigilance, and the moment-to-moment fluctuations of this arousal change can be tracked using the HAS and LAS. In addition, PVT performance was found to have a positive correlation with the proportion of occurrence in the HAS, and negatively correlated to the proportion of LAS. Second, an individual’s proportional preponderance of these two FC states, as combined into the AI, can be used to predict subsequent vigilance declines over five nights of sleep restriction (Patanaik et al. 2018).

The current experiment builds on these results by showing that fluctuations in HAS/LAS proportions are correlated with SD vulnerability when sleep pressure is manipulated experimentally, strengthening the case that these specific states robustly index levels of vigilance. This is important, as a variable that predicts a future outcome may not necessarily be the same variable that changes when that outcome is realized.

The interest in individual differences in SD vulnerability originates from behavioural observation that, over multiple nights of SD, declines in vigilance are stable within individuals but highly variable between them (Van Dongen et al. 2004). Hypoactivation in dorsal attention areas is also stable over multiple SD nights (Lim et al. 2007), suggesting that fMRI may effectively capture this vulnerability. Supporting this, fMRI activity in frontoparietal regions, visual cortex, and the thalamus is attenuated during PVT lapses following SD, as compared to behaviourally similar lapses following normal sleep (Chee et al. 2008). In that experiment, slower responses during SD elicited lower activity in both intraperietal sulcus and inferior occipital cortex, whereas lower activity was found only in inferior occipital cortex for faster responses.

These results notwithstanding, there is still a lack of evidence directly linking brain activity to individual differences in SD vulnerability in the domain of vigilance, which is most substantially affected by acute sleep deprivation: most research to date has focused on selective or orienting attention (Ma et al. 2015). Our current findings provide some data to bridge that knowledge gap, robustly linking two DCSs to this trait-like phenomenon.

While it may appear contradictory that %HAS correlates with SD vulnerability without significantly decreasing in the SD state, this is in fact reinforces the idea that some proportion of HAS is essential to maintain engagement with the external environment. In other words, declines in vigilance might be necessary, but not sufficient to cause significant reductions in HAS. Future research might investigate longer durations of SD to interrogate whether significant HAS declines occur in parallel with more serious cognitive failure (e.g. PVT “timeouts” of > 30 s).

DCS-vigilance correlations were not observed in any of the non-arousal related states, even though their proportions changed significantly following SD. Of the three remaining non-arousal states, we have previously described the TRS as being related to trait mindfulness (Baer et al. 2008): individuals who scored higher on a test of objective mindfulness spent more time in this state (Lim et al. 2018). Interestingly, while mindful individuals also have greater attentional capacity and show superior performance on the PVT (Wong et al. 2018), this association is not seen in the current dataset, in which the PVT declines over SD are driven by decreases in arousal and not mindfulness. This dissociation is further evidence supporting the specificity and sensitivity of our named DCSs. Finally, declines in self-reported mindfulness have been reported following multiple-day sleep restriction (Campbell et al. 2018), and this is in line with our observation of significant decreases in TRS following SD. However, the lack of mindfulness measures with our current cohort renders this explanation speculative.

### Decrease in state transitions after sleep deprivation are also associated with behaviour

In exploratory analysis, we found that change in PVT performance across state was correlated with the change in the percentage of state transitions during resting-state scans. We have previously proposed a link between state transitions and the ability of the brain to refocus attention (Lim et al. 2018), which may represent a marker of cognitive flexibility (Li et al. 2017, Marusak et al. 2018); this theory is supported by data in macaques showing that sedation is associated with a loss of the rich repertoire of states seen in wakefulness (Barttfeld et al. 2015). The negative effects of sleep deprivation on cognitive flexibility are well known (Harrison and Horne 1999, Durmer and Dinges 2005), with increasing sleep pressure interfering with top-down executive function maintained by the prefrontal cortex (PFC). In addition to PFC dysfunction, SD is associated with more variable cognitive performance as the top-down drive to remain vigilant competes with the homeostatic drive to fall asleep (Doran et al. 2001, Goel et al. 2009), a phenomenon that has been termed *wake-state instability*.

Wake-state instability predicts that more frequent transitions would be observed in the connectivity time course following SD, reflecting more rapid switches between dorsally generated RW-associated states and centrally generated sleep-promoting ones. For example, in an fMRI study of attentional lapsing, Chee et al. (2008) showed that periods of fronto-parietal hypoactivation and thalamic compensation after SD were interspersed among trials that were comparable with the rested state. Unexpectedly, this was not what we observed in the current dataset – transitions between DCSs decreased in the SD state. One possible reason for this finding is that the moving average in our DFC analysis smoothed over the more abrupt state transitions associated with wake-state instability, and the remaining decreases in DCS transitions more exclusively reflect top-down executive failure to refocus attention. A more plausible explanation may be that unstable brain dynamics are only observable when a participant is challenged with a task and not in an unconstrained resting-state scan in an environment that favours falling asleep.

### No additional predictive information from other physiological metrics to individual differences in vigilance decline

Aside from our primary analysis, we interrogated two other variables known to change after SD to control for these potential confounds.

The relationship between global signal (GS) and vigilance has been the subject of numerous studies (Wong et al. 2013), and has been found to be predictive of vulnerability to SD (Patanaik et al. 2018). Our results suggest that there might be a more nuanced relationship between GS, DCS, and arousal. While we found a significant univariate correlation between GS and with change in lapses, GS did not contribute a significant incremental effect to predicting SD vulnerability when modelled together with AI. In contrast, prior work found that both global signal and AI as a whole predicted changes in vigilance decline (Patanaik et al. 2018). Since GS fluctuations have previously been associated with transitions between states of varying arousal (Liu et al. 2018), we propose that our null findings may be due to significant overlaps between the contributions of GS and AI. In a similar vein, head motion is also found to have a null contribution to the relationship between AI, vigilance, and GS. Given that GS and head motion are closely related as well (see Laumann et al. 2017), the lack of independent contribution from head motion is unsurprising.

## Conclusion

In summary, we have established that total sleep deprivation affects the occurrence of specific DCSs that may relate to arousal. Converging evidence from several studies suggests that these DCSs are reproducible, and can meaningfully be used as an index of arousal to track changes in vigilance following total sleep deprivation. Such coupling between changes in arousal-related DCSs and behavioural changes persist beyond physiological interferences such as head motion and the global signal. Further, these DCSs have been found in multiple studies employing different methodologies, and their similarities suggest the robustness of these specific states in their predictive utility.

## Acknowledgements

This work was supported by the National Medical Research Council, Singapore (STaR/0015/2013). We acknowledge the assistance of Vinod Shanmugam in data collection.

